# EEG-based diagnostics of the auditory system using cochlear implant electrodes as sensors

**DOI:** 10.1101/2020.07.16.206250

**Authors:** Ben Somers, Christopher J. Long, Tom Francart

## Abstract

The cochlear implant is one of the most successful medical prostheses, allowing deaf and severely hearing-impaired persons to hear again by electrically stimulating the auditory nerve. A trained audiologist adjusts the stimulation settings for good speech understanding, known as “fitting” the implant. This process is based on subjective feedback from the user, making it time-consuming and challenging, especially in paediatric or communication-impaired populations. Furthermore, fittings only happen during infrequent sessions at a clinic, and therefore cannot take into account variable factors that affect the user’s hearing, such as physiological changes and different listening environments. Objective audiometry, in which brain responses evoked by auditory stimulation are collected and analysed, removes the need for active patient participation. However, recording of brain responses still requires expensive equipment that is cumbersome to use. An elegant solution is to record the neural signals using the implant itself. We demonstrate for the first time the recording of continuous electroencephalographic (EEG) signals from the implanted intracochlear electrode array in human subjects, using auditory evoked potentials originating from different brain regions. Furthermore, we show that the response morphologies and amplitudes depend crucially on the recording electrode configuration. The integration of an EEG system into cochlear implants paves the way towards chronic neuro-monitoring of hearing-impaired patients in their everyday environment, and neuro-steered hearing prostheses, which can autonomously adjust their output based on neural feedback.

## I. Introduction

The cochlear implant (CI) is one of the most successful medical prostheses, allowing deaf or severely hearing-impaired persons to regain their hearing. The CI system consists of an internal part implanted during a surgery, and an external part worn behind the ear [Dorman and Wilson, 2004; Zeng et al., 2008]. Sound is captured with a microphone and transformed by the speech processor into a sequence of electrical stimulation pulses. This sequence is encoded and transmitted to the implant through a wireless radio-frequency link. The electrical stimulation is applied to the implant’s electrode array located in the cochlea, stimulating the auditory nerve and causing audible percepts for the user.

Despite the success of cochlear implants, there is a large variability among users’ hearing outcomes such as speech intelligibility, especially in noisy listening environments. One factor contributing to this variability is the difficulty of finding the optimal sound processing and stimulation settings for an individual user. The transformation of an acoustic signal to an electrical pulse sequence involves the tuning of hundreds of parameters, referred to as “fitting” the cochlear implant [Wouters et al., 2015]. Fitting is performed by specialized audiologists during infrequent sessions in the clinic, and often relies on behavioral feedback from the CI user. This makes the procedure time-consuming and difficult, especially for people who cannot actively collaborate such as deaf children or persons with a disability. Due to time constraints, the fitting procedure generally starts from a default setting, of which only a subset of parameters is varied based on the behavioural feedback from the user. Furthermore, due to changes in physiology, anatomy, and subjective preferences over time, the fitting might ideally be updated frequently. However, for the majority of of CI users, the settings are updated at most yearly or even less frequently.

Improvements to the current CI fitting paradigm are expected from the field of objective audiometry, in which neural responses evoked by auditory stimulation are collected. An advantage of the evoked response approach is that active participation of the user is no longer required, which eliminates confounding factors such as motivation, attention, and cognitive state. Electroencephalography (EEG) is an essential tool to acquire such brain responses due to its high temporal resolution and relatively low cost and complexity. As such, recent years have seen a lot of research involving the estimation of EEG-based objective measures relevant for the fitting of hearing devices (e.g. estimating auditory thresholds [Brown et al., 2000; Campbell et al., 2016; Koka et al., 2017], comfortable loudness level and loudness growth [Brown et al., 2000; Botros and Psarros, 2010; Visram et al., 2015; Van Eeckhoutte et al., 2018], neural survival in cochlea [Kang et al., 2018], …).

Unfortunately, the evoked potentials (EP) currently in use in clinical practice only target the peripheral components of the hierarchical auditory pathway. For instance, the electrically evoked compound action potential (ECAP) measures the synchronous firing of a population of stimulated auditory nerve fibres [He et al., 2017]. This response occurs within one millisecond post-stimulus and represents the first stage of stimulus encoding on the auditory nerve. The auditory brainstem response (ABR) is another short-latency evoked potential, originating from the brainstem within a 10 millisecond post-stimulus window, which can be useful for estimation of auditory thresholds [Truy et al., 1998; Brown et al., 2000; Kubo et al., 2001]. Relevant measures for high-level functioning of the auditory system such as speech intelligibility originate from cortical brain regions and typically occur at much larger latencies than responses from the periphery. Recording of long-latency responses such as cortical evoked potentials (CEP) requires recording windows up to hundreds of milliseconds. Furthermore, auditory measures in response to continuously ongoing stimuli have been shown to relate to speech intelligibility, such as the auditory steady state response (ASSR) [Gransier et al., 2019] or speech envelope tracking responses [Ding and Simon, 2013; Vanthornhout et al., 2018; Somers et al., 2018; Lesenfants et al., 2019; Verschueren et al., 2019]. Measurement and analysis of these responses requires continuous EEG recordings.

In summary, EEG-based measures show great potential for the development of better, objective, and in the future even automatic fitting procedures. However, recording EEG still requires considerable time and effort in the clinic. For optimal signal quality, the skin should be abraded and a conductive gel must be applied before placing the electrodes on the scalp. Furthermore, participants are required to remain still for the duration of the recording session to avoid motion artifacts contaminating the obtained data, making EEG recordings generally an unpleasant experience for the subject. Another issue is that the soundproof EEG rooms in the clinic are not a realistic listening environment which a CI user may face in their everyday life. Mobile EEG recording systems which can be autonomously applied and worn by the CI user can solve some of these issues. Several wireless EEG systems have appeared on the market, but these are still too heavy and too bulky for frequent and unobtrusive and neuro-monitoring.

An elegant solution is to use the implanted electrodes of the CI. Current commercial CIs have some basic telemetry capabilities, but are limited by hardware constraints of the embedded electronics. The recording amplifiers are limited to very short recording windows, which can only capture measures of the auditory periphery [Campbell et al., 2015; Abbas et al., 2017; Tejani et al., 2019]. By repeating the measurement multiple times with a recording window shifted to larger post-stimulus latencies and combining all of these recording windows together, longer-latency responses can be recovered [Mc Laughlin et al., 2012; Abbas et al., 2017]. However, due to the repeated measurements, this procedure drastically increases recording time. Furthermore, the clinical software used in these studies is not designed for such recordings, requiring manual adjustment of recording parameters for every repetition. This renders the method of stitching together multiple recordings impractical and time-consuming. Some studies have demonstrated the recording of neural responses to electrical stimulation using (non-cochlear) implanted electrodes, for instance with a few epidural electrodes [Haumann et al., 2019] or an intracranial grid [Nourski et al., 2013] placed during surgery. These methods are highly invasive and only appropriate for temporary use.

In this study, we demonstrate the recording of continuous EEG from the implanted electrodes of a cochlear implant. We recruited a special population of CI users with a percutaneous connector, allowing direct access to the implanted electrodes. To the authors’ knowledge, continuous EEG has never been recorded before with CI electrodes in humans. In this study, we targeted two well-established evoked responses: the electrically evoked ABR (EABR) and the electrically evoked CEP (ECEP), respectively a short-latency subcortical response and a long-latency cortical response. These experiments offer the unique opportunity to explore the characteristics of cochlear implant EEG recordings and the impact on evoked responses, such as the optimal implanted electrode configuration for EEG-based objective measures, the manifestation of the electrical artifact, and possible differences with recordings using conventional scalp electrodes. These are important characteristics to consider for the future integration of EEG systems into cochlear implants, and may guide design decisions for optimal measurement of objective measures using CIs. Potential applications of EEG-CI systems are chronic neuro-monitoring of CI users, remote fitting and diagnostics, and so-called neuro-steered hearing devices [Van Eyndhoven et al., 2016; Geirnaert et al., 2019; Aroudi and Doclo, 2020], which autonomously adapt their fitting based on responses from the user’s brain.

## II. Results

Electrically evoked auditory potentials were recorded in three subjects with an experimental CI device provided by Cochlear Ltd., which allows direct access to the implanted electrodes through a percutaneous connector. The locations of the different electrodes used in the experiments are indicated in Fig. 1. The implanted electrodes are either part of the 22-electrode array placed in the cochlea, or two extracochlear reference electrodes. The intracochlear electrodes are numbered E01 to E22, counting from base towards the apex of the cochlea. The extracochlear electrodes are referred to as MP1 and MP2 (i.e. monopolar electrodes 1 and 2). The MP1 electrode is a small ball electrode placed under the temporal muscle. In conventional CIs from Cochlear, the MP2 electrode is a plate electrode attached to the casing of the implant. In the percutaneous devices used in this study, there is no implant casing, so the MP2 electrode is a flat plate electrode also placed under the temporal muscle. Four Ag/AgCl electrodes were placed on the scalp in locations commonly used in evoked potential paradigms.

**Fig. 1.**
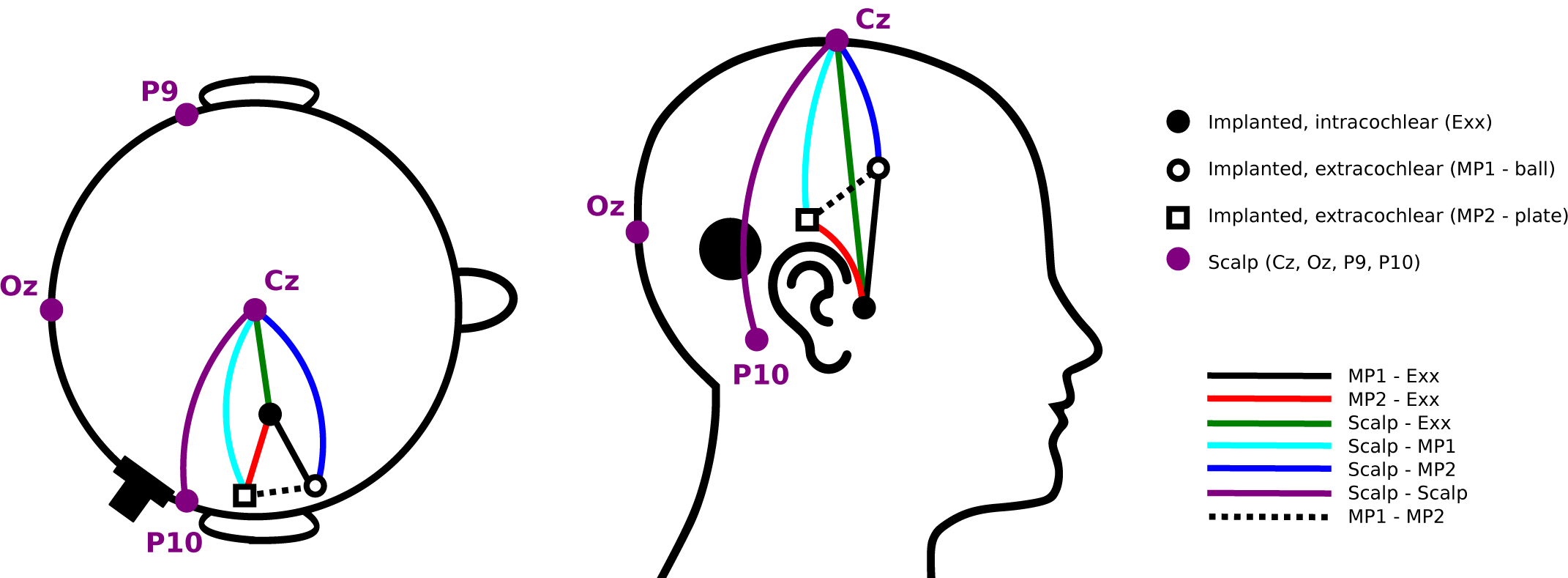
Overview of locations of the electrodes used for recording evoked potentials (not to scale). Implanted electrodes are either located in the electrode array in the cochlea (numbered E01 to E22), or are one of the two extracochlear reference electrodes located in the temporal muscle (MP1 and MP2). Scalp electrodes are placed in central (Cz), occipital (Oz) and left/right mastoid (P9/P10) sites. In this instance, the percutaneous connector is shown on the right side of the head. The different recording dipoles between which EEG signals are measured are indicated with color-coded lines between electrodes. This color-coding scheme is used for indicating recording configurations of evoked potentials in subsequent figures.

Our experimental set-up allowed to record EEG signals between any two electrodes. We report evoked potentials recorded between extracochlear and intracochlear implanted electrodes, i.e. MP1-to-Exx or MP2-to-Exx (where Exx is a numbered intracochlear electrode). For comparison with the conventional, clinical EP paradigm, we also recorded between two scalp electrodes. Finally, we obtained recordings between one implanted and one scalp electrode, which we will refer to as “hybrid” recordings as opposed to the implanted and scalp recordings. The hybrid recordings have the advantage that the scalp electrode can be moved around freely. This allows to use the scalp electrode to determine the recording configuration that optimally captures the EP dipole. This information can guide placement of implantable recording electrodes in future integrated CI-EEG systems. The color-coding of recording configurations in Fig. 1 is reused in subsequent figures.

The targeted neural responses were evoked using biphasic electrical pulse trains with 100 µs phase duration and an inter-phase gap of 8 µs. All stimulation was presented in bipolar mode, i.e. between two intracochlear electrodes, such that both extracochlear reference electrodes were available for recordings. EABRs were evoked using a single biphasic pulse with a repeat frequency of 27 Hz. ECEPs were evoked using a burst of 10 biphasic pulses during 5 ms with a repeat frequency of 1 Hz. Further stimulus details can be found in Section IV-D.

### A. Electrically evoked auditory brainstem responses - Intensity sweeps

Fig. 2 shows EABR intensity sweeps obtained for all three subjects. During a sweep, EABRs are obtained using stimuli with intensities ranging from the loudest comfortable level (C-level) to below the audible threshold level (T-level). The stimulus pulse occurred at 0 ms, followed by a 1 ms blanking window during which the recording amplifier gain was set to zero to prevent amplification of the electrical stimulus artifact [Kent et al., 2015]. The EABR response was reliably recorded for all subjects and is visible in the 1 ms to 5 ms window. Simultaneous recordings using either MP1 or MP2 as active electrode are shown, with an intracochlear reference electrode, which was chosen depending on the subject.

**Fig. 2.**
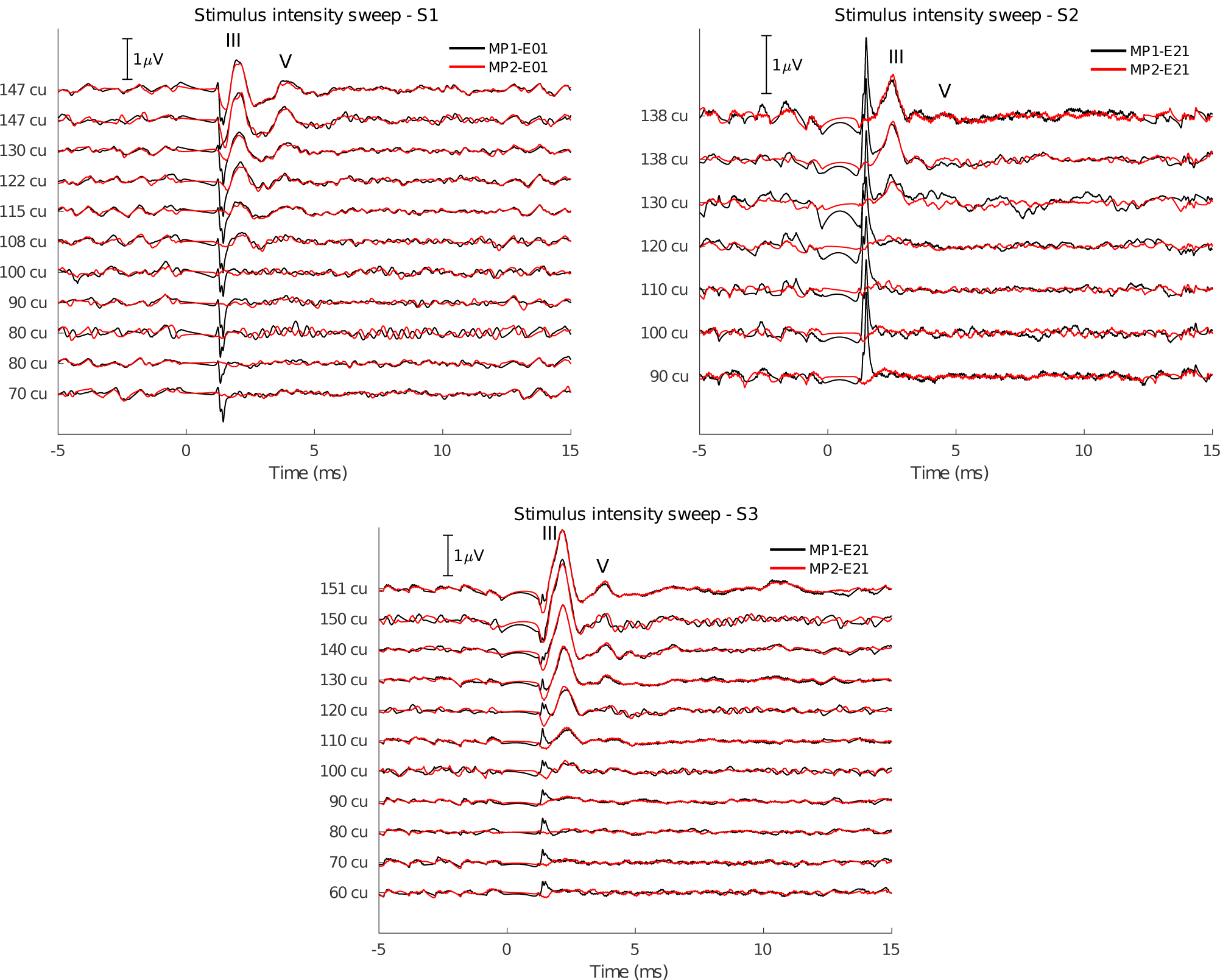
EABRs recorded at varying stimulus intensities. Waves III and V are marked on the traces. The recording at the loudest comfortable level was repeated twice. Electrical stimulation through the CI was bipolar using electrodes E21-E13 for S1 and E04-E12 for S2 and S3. The intracochlear recording electrode depends on the subject (S1: E01, S2 and S3: E21). Stimulation intensity is given in current units (cu), a logarithmic function of current (in µA) used to account for the non-linear loudness growth in the human auditory system.

The most prominent components of the EABR are wave III, occurring at a latency of 2 ms, and wave V, occurring at a latency of 4 ms. Waves III and V are reliably recorded for all subjects, except for subject S2, whose wave V is almost absent from the C-level recordings. As expected, the EABR amplitudes gradually decrease for lower current levels. Additionally, the latency of wave V increases for lower current levels. Interestingly, wave III is larger than wave V in all recordings. This is opposite to scalp recordings, where wave V is usually the largest and most distinguishable wave of the response complex. This can be explained by the closer proximity of the cochlear recording electrodes to the neural generator of wave III, whereas the generator of wave V is located further along the auditory pathway in the brainstem.

There are no differences in EABR amplitudes and latencies between recordings using MP1 or MP2 as active recording electrode. The MP1 recordings consistently contain an artifact at the offset of the blanking (at 1 ms). It is a non-neural signal as it does decrease in amplitude for lower current levels, even sub-threshold. This artifact may be explained by the different shapes of electrodes MP1 and MP2: electric fields are generally stronger near sharper surfaces with a higher curvature [Liu, 1987; McAllister, 1990]. This may cause the smaller MP1 ball electrode to be more sensitive to residual stimulation artifact or electrical artifacts caused by switch-off of the amplifier blanking compared to the larger MP2 plate electrode. Additionally, despite MP1 and MP2 being in similar locations, differences in the electrical characteristics of the surrounding tissue, such as skin and bone, can contribute to different discharge dynamics [Freche et al., 2018].

### B. Electrically evoked auditory brainstem responses - Electrode configurations

The results of the intensity sweep (Fig. 2) show that the EABR response is present for all subjects at their C-level, measured entirely with implanted electrodes. In the following experiment, the recording electrodes are changed to obtain different configurations, including scalp electrodes, to assess changes in EABR morphology and determine the optimal recording configuration.

Fig. 3 shows EABR responses recorded with different recording electrode configurations using stimulation at C-level for all three subjects. The MP1-to-intracochlear and MP2-to-intracochlear recordings from Fig. 2 are shown again for reference. Due to timing constraints, not all configurations were tested for all subjects. In some configurations, the recording was repeated, indicated with multiple overlaid traces.

**Fig. 3.**
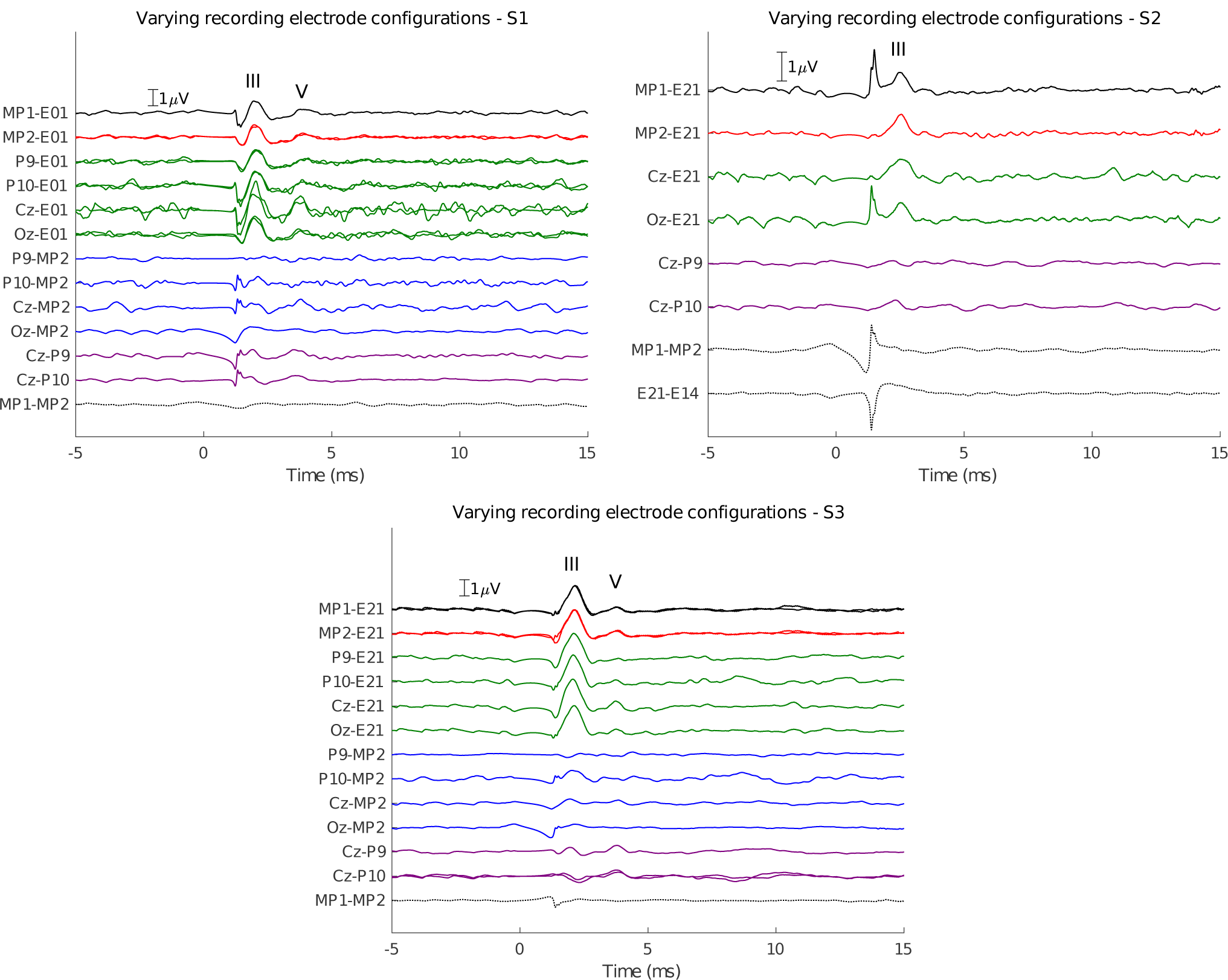
EABRs recorded using various electrode configurations, indicated on the vertical axis and color-coded according to Fig. 1. Waves III and V are marked on the traces. Repeated tests with the same recording electrode configuration are overlaid. Electrical stimulation through the CI was bipolar using electrodes E21-E13 for S1 and E04-E12 for S2 and S3.

Generally, the EABR response is also observable from hybrid (i.e. scalp-to-implanted electrode) recordings when an intracochlear recording reference electrode is used (green traces). In these hybrid recordings, wave III is always the most distinguishable wave in the response. Wave V is most pronounced for the hybrid recordings using the Cz (central) electrode. This is interesting, because Cz-to-Exx most closely resembles the usual clinical montage for EABR recordings, i.e. Cz-to-mastoid. This implies that an implanted reference electrode closer towards the top of the head would result in better EABR recordings for wave V.

For hybrid recordings with an extracochlear reference electrode, the EABR is difficult to detect. This was only tested for S1 and S3 using scalp-to-MP2 recordings. The wave III is only present when using the P10 and Cz scalp electrodes, albeit smaller compared to the intracochlear reference electrode case. Wave V is small and only present for the Cz configuration. The small amplitudes (or absence) of these responses can be explained by the orientation of the recording dipoles formed between scalp electrodes and MP2. Note that both S1 and S3 had the CI in the left ear, so P10 is contralateral to the MP2 electrode and therefore captures a larger dipole.

For S1 and S3, the pure scalp recordings (Cz-P9 and Cz-P10) show normal EABRs with distinguishable waves III and V. Wave III is much larger when recorded from implanted electrodes compared to scalp electrodes. For S2, the EABR is not present in scalp recordings. Subject S2 is likely to have a small, difficult to detect EABR response, which is also evidenced by the small EABR at comfort level for this subject in Fig. 2.

We did not detect EABR responses from MP1-to-MP2 recordings in any subject. Likely, these two reference electrodes are too close together and in a plane that does not capture the EABR response dipoles well. Similarly, the E21-to-E14 recording recording between two intracochlear electrodes in S2 did not lead to a response because both electrodes are too close together relative to the distance of the cochlea to the generator of the EABR.

### C. Electrically evoked auditory cortical responses - Electrode configurations

The cortical response complex is characterized by P1 (50 ms) - N1 (80-100 ms) - P2 (150-180 ms) - N2 (200-250 ms), of which the N1 is usually most pronounced as a negative deflection. This response was recorded for all three subjects with different recording configurations, shown in Fig. 4. The responses were evoked using short bursts of electrical pulses every 1 second at C-level.

**Fig. 4.**
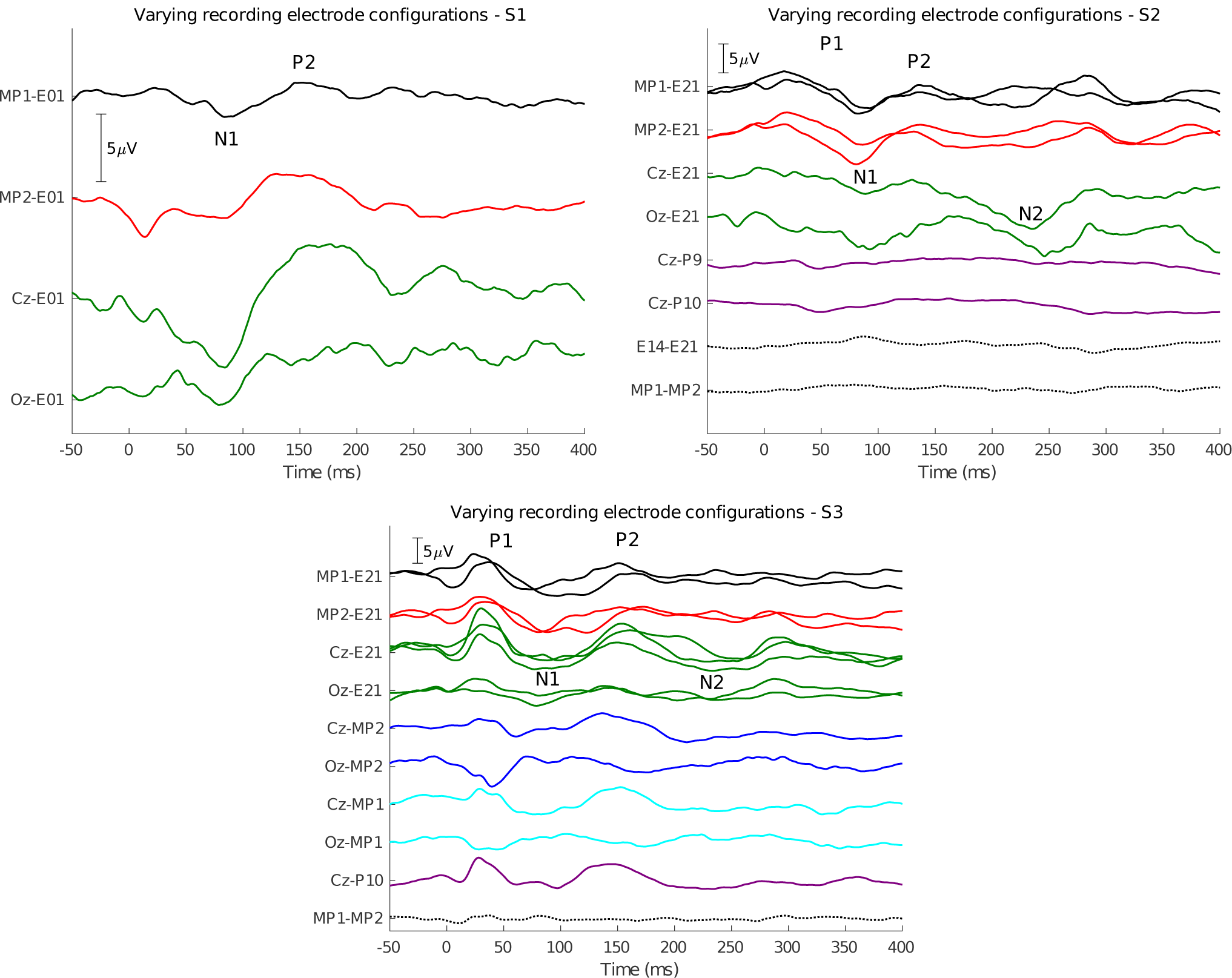
ECEPs recorded using various electrode configurations, indicated on the vertical axis and color-coded according to Fig. 1. P1, N1, P2 and N2 are marked on the traces where applicable. Repeated tests with the same recording electrode configuration are overlaid. Electrical stimulation through the CI was bipolar using electrodes E21-E13 for S1 and E04-E12 for S2 and S3.

The different components of this response can be observed from the fully implanted electrode recordings, but are most clear when using a Cz scalp electrode in a hybrid recording configuration. Similarly to the results for the EABR, this configuration resembles the Cz-to-mastoid montage that is most often used for recording cortical responses.

For subject S1, due to timing limitations, only a few configurations were recorded. The N1-P2 components are clear, especially with the favourable Cz-E01 montage. For subject S2, the cortical response was found when using implanted or hybrid recording montages, but not when using solely scalp electrodes. Similarly to the EABR in this subject, the responses are likely very small and thus difficult to detect from noisy scalp recordings. Additionally, implanted electrode recordings using pairs of extracochlear or pairs of intracochlear electrodes did not work. For subject S3, very clear P1-N1-P2-N2 complexes are observed, which are also reliable across repetitions, especially for the hybrid Cz-to-E21 recording. The response is also present from hybrid recordings between Cz and MP1 or MP2 as the reference electrode, but much less when Oz is used as the active electrode. This again marks the importance of using a recording electrode in a favourable (in this case, central) recording site. The peak-to-peak ECEP amplitudes are larger when using the Cz-to-E21 recording pair compared to Cz-to-MP1 or Cz-to-MP2 (only available for S3), which implies an advantage of using an intracochlear recording reference electrode compared to an extracochlear one, i.e. this results in lower electric field attenuation by the skull. For S3, the scalp electrodes also show a clear cortical response. No responses were obtained from the extracochlear (MP1-to-MP2) recording.

### D. Completely internal recordings

In the experiments and results shown so far, for simplicity’s sake an electrode on the subject’s collarbone was used as the ground for all EEG recordings (see section IV-C). In a future integrated CI-EEG system, there will likely be no percutaneous wires. The EEG recording system should be completely implanted, and an external ground electrode would be impossible. We explored whether using an implanted ground electrode impacts the measured responses in an additional condition with subject S2. Fig. 5 compares the EABRs recorded from implanted electrodes. In the figure, either the collarbone electrode or the other MP electrode was used as recording ground. The recorded EABR does not show any meaningful differences. However, the artifact at 1 ms picked up by the MP1 electrode is absent when using MP2 as a ground. The MP2 ground electrode likely provides a nearby, low-impedance path for charges to quickly dissipate, which reduces the sensitivity of the MP1 electrode to electrical artifacts [Freche et al., 2018].

**Fig. 5.**
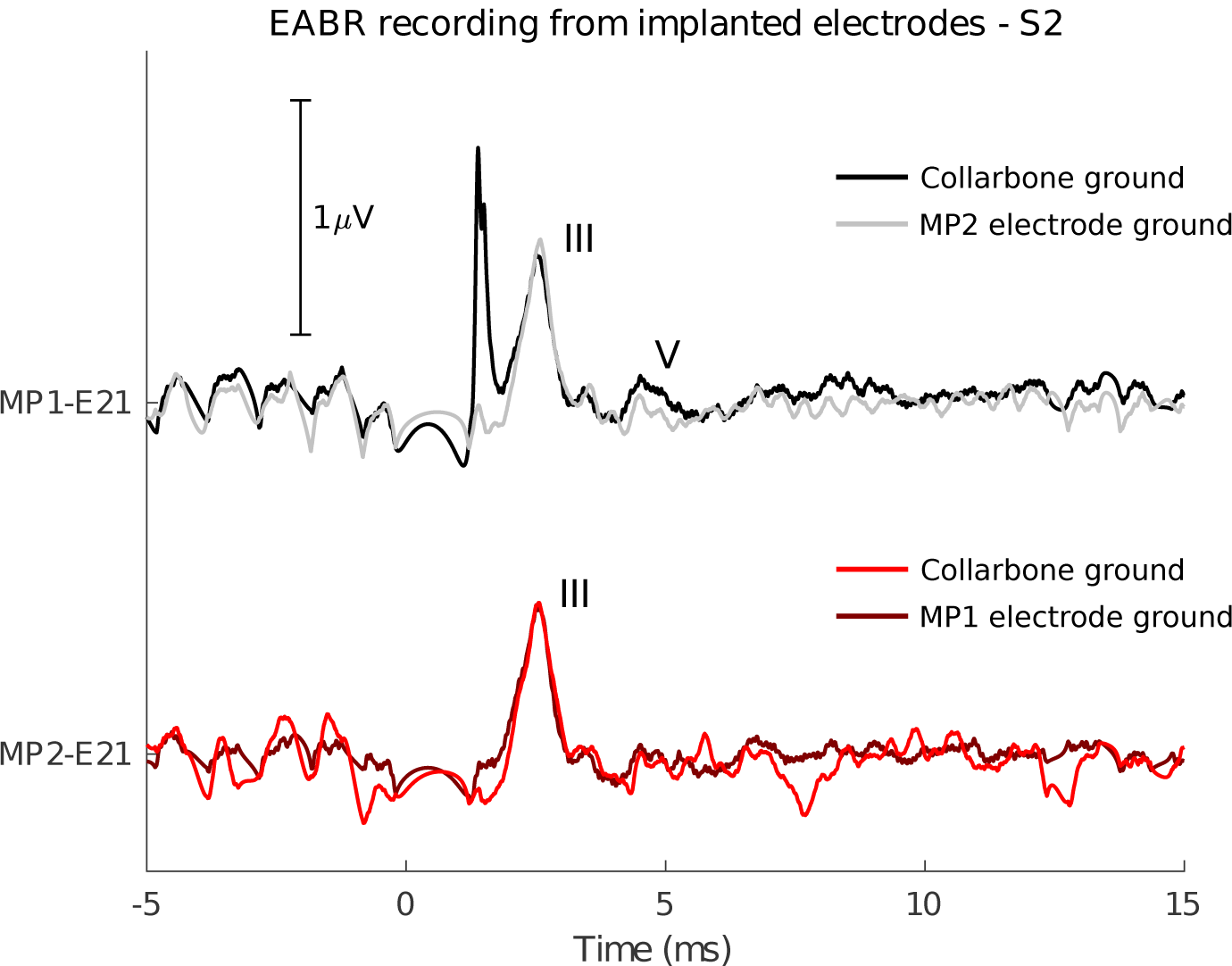
Comparison between EABRs using an external ground electrode (collarbone) or an implanted ground electrode (MPx). Stimulation settings were identical to those used for other EABRs obtained at C-level.

## III. Discussion

EP measurements using the implanted electrodes of a CI are available in the devices of all major CI manufacturers. However, due to hardware limitations of the embedded implanted electronics, only short recording windows of a few milliseconds can be buffered and transmitted to the external part. This only suffices to capture very low-latency responses (e.g. ECAPs) from the the auditory nerve [Abbas et al., 1999; Brown et al., 2000; Tejani et al., 2019]. Mc Laughlin et al. [2012] developed a technique to record EEG from cochlear implant electrodes using the built-in telemetry functionality of a CI from Cochlear Ltd. The clinical implementation of the forward masking paradigm [Brown et al., 1990] in the Custom Sound EP software was used, which provides recording windows limited to 1.6 ms. The targeted longer-latency responses were obtained by patching together multiple overlapping recording windows obtained at different latencies, which required a lot of stimulus repetitions. Recording time was increased even more because the forward masking paradigm requires four stimulus repetitions with different masker and probe pulse configurations per recording window. The authors reported that data collection was time-consuming and cumbersome, causing multiple stimulus conditions to be shortened. For longer-latency EPs like the ECEPs, only a few data points at latencies where peaks were expected were obtained. In our study, we have complete control over the amplifier and recording parameters, allowing us to measure continuous EEG in a straightforward way without the need for stimulus repetition.

Mc Laughlin et al. [2012] only partially recorded EABR responses from CI electrodes at three stimulus intensity levels. They found that wave V had an amplitude comparable to scalp recordings, but a larger wave III. This was hypothesized to have two plausible causes: firstly, the stimulation artifact was not accounted for in their EABR recordings, causing earlier waves of the EABR to be elevated. Secondly, the intracochlear electrodes are situated closer to generators of early EABR components, such as the auditory nerve and brainstem, causing the electric potential at the electrodes to be larger compared to scalp electrodes. Our results confirm this observation (Fig. 3), but since we accounted for the the stimulus artifact in various ways, we can explain the difference by the closer proximity to the neural generator. We were also able to record the cortical response complex from intracochlear recordings, even for subjects where the response was not clear from scalp recordings (e.g. S2 on Fig. 4). As we had the freedom to move around a scalp electrode used as the active recording electrode, we showed that the recorded EP can even be larger if the recording electrode is more appropriately placed to capture the neural response dipole (i.e. more towards the central/occipital area).

Several studies have employed other techniques for invasive recordings of auditory evoked potentials in human subjects without using CI electrodes. For longer-latency neural responses originating from higher brain regions, a major hurdle is that subjects generally need to be conscious. For instance, during surgery under general anaesthesia, such as cochlear implantation, only EPs originating from the auditory periphery (i.e. auditory nerve, brainstem) can be elicited, but cortical responses can not [Heinke and Koelsch, 2005; Nourski et al., 2013]. Early responses such as the EABR are useful as they reflect low-level neural processing of sound in the brain, but they do not correlate well with higher-level auditory functions such as speech understanding. Haumann et al. [2019] placed epidural electrodes over the auditory cortex during an otherwise conventional CI surgery. Three extra holes were drilled through the skull down to the dura. In the days following surgery, the percutaneous electrode leads were connected to an external EEG amplifier for several evoked potential recordings. The epidural electrodes were removed after the recording sessions, 4-5 days post-surgery. For their EABR recordings, they did not find an improvement when recording with epidural electrodes compared to scalp electrodes. However, the epidural electrode were positioned over the auditory cortex to optimize recording of ECEPs. For cortical responses, the authors report clearer cortical responses compared to scalp electrodes which are visible at lower stimulation intensities and less affected by artifacts compared to scalp electrodes. Our results show equal (S3) or better (S2) ECEP responses recorded with implanted compared to scalp electrodes, and even larger ECEP responses when optimizing the electrode positions through hybrid (Cz-to-implanted) recordings. Nourski et al. [2013] recorded auditory EPs in a single CI user using a subdural electrode grid which was placed as part of clinical epilepsy treatment. The authors used these intracranial electrophysiological recordings to study the response properties of a region of the non-primary auditory cortex. These studies were highly invasive as they required drilling of extra holes through the cranium, and implantation of additional recording equipment.

### Integration of EEG systems into CIs

We have shown that it is possible to use the existing electrodes of a CI to record continuous EEG for both short and long-latency EPs. By dedicating one electrode of the intracochlear array and one extracochlear reference electrode, current CIs could be used for EEG recordings without changing the electrodes, provided some upgrades to the embedded electronics on the implant that would allow to buffer, compress and transmit the data. Note that CIs from Cochlear Ltd. have two extracochlear reference electrodes, i.e. the MP1 ball electrode placed under the temporalis muscle and the MP2 plate electrode on the implant casing [Cohen et al., 2002]. Devices from other manufacturers, such as Med-El or Advanced Bionics have one extracochlear electrode located on the implant casing. This electrode is usually reserved for stimulation and thus cannot be used for simultaneous recording. Even for Cochlear CIs, both MP1 and MP2 electrodes are often used in parallel as return electrodes for clinical stimulation schemes, referred to as the MP1+2 configuration.

Another argument for the inclusion of dedicated recording electrodes is that the current placement of the reference electrode (either casing or ball) is not optimal for EEG recordings. Our experiments with hybrid recording configurations reveal differences in response amplitude across different scalp electrode locations, caused by the orientation of the recording electrode pair relative to the EP dipole in the brain. These relative amplitude differences between recording sites will still hold if the scalp electrode would be implanted subcutaneously similar to the MP1 reference electrode, or even intracranial similar to electrocorticographical (ECoG) electrodes used in BCI applications [Leuthardt et al., 2004]. Intracranial electrodes will record even larger responses as the skull greatly attenuates electric fields generated in the brain [Schalk and Leuthardt, 2011]. This reduces the required gain for amplifying the signals and thus saves power. However, this would come at the cost of increased invasiveness of the surgery.

Ideally, implanted reference electrodes for either stimulation or recording should be placed in different locations. The stimulation reference is usually situated near the implant casing under the temporalis muscle: the further away it is placed from the intracochlear electrodes, the higher the power consumption while stimulating [Ramos-Miguel et al., 2015]. An electrode dedicated to recording EEG should be implanted close to the central (Cz) area if the other electrode of the recording pair is located near the ear, such as an intracochlear electrode, an extracochlear electrode under the temporalis muscle, or an electrode near the mastoid. As demonstrated in our experiments, having one electrode of the recording pair in the cochlea is favourable for capturing larger brain responses, as they are less attenuated by the skull and less susceptible to susceptibility to external noise and artifacts.

### Future outlook: closed-loop CI systems and neuro-steered hearing devices

The capability of a CI system to record neural feedback, in an unobtrusive way and without requiring additional recording equipment, is a building block for several envisioned applications. This technology can be used for making the CI a closed-loop system [Wilson et al., 2011; Mc Laughlin et al., 2012]. In the current rehabilitation paradigm, CI stimulation settings are only adjusted during infrequent clinical fitting sessions, in between which the device operates as an open loop system (Fig. 6A). It does not incorporate feedback from the user, despite several factors that vary over time and may prompt a change in the CI’s settings, such as progression of the hearing loss, different listening environments, or attentional state of the user. These factors can be used to adapt audio processing settings in the short term (e.g. gain control, noise suppression) or the fitting in the long term (e.g. threshold and comfortable levels, pulse rate, pulse shape and width).

**Fig. 6.**
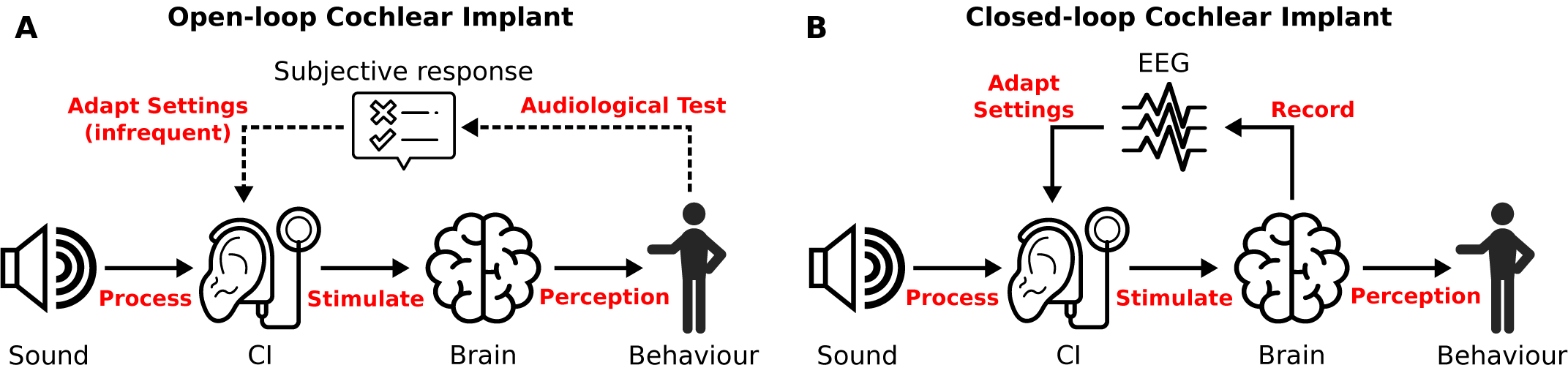
**A** Currently, CIs operate as an open loop system for the majority of the time. Behavioural feedback from the user is used to fit the device settings during infrequent visits to an audiologist. **B** Neural feedback can be used to close the loop. This eliminates the need for behavioural feedback, and can happen continuously if an EEG system is integrated with the CI. *Image composed with icons from thenounproject*.*com, see acknowledgements*.

In a first realization, neural responses can be monitored by the device to be logged and transmitted to an audiologist, who can remotely assess the diagnostics. If required, the audiologist can invite the user for a fitting session in the clinic, or even perform a remote fitting procedure. One step further is to have devices perform the fittings themselves based on the objective auditory diagnostics [Mc Laughlin et al., 2012; Visram et al., 2015; Finke et al., 2017], which can be collected intermittently or continuously (Fig. 6B). An additional advantage of such a closed-loop CI is that the auditory diagnostics are obtained using natural sound and speech from the user’s everyday environment, whereas clinical audiological tests often make use of specifically crafted, non-natural speech materials. Furthermore, objective EEG-based diagnostics don’t require active participation from the user. This technology can be applied to better and more efficiently fit the CIs of difficult-to-test populations, such as young children, or persons with a communication impairment.

Various types of evoked potentials are useful diagnostics for the auditory system, and usually correlate with low-level functioning of the auditory pathway, such as neural survival in the cochlea, hearing thresholds and comfortable loudness levels. However, with the possibility to record continuous EEG, measures that correlate with higher-level aspects of hearing can be used to steer hearing devices. An EEG-based measure that predicts speech understanding [Vanthornhout et al., 2018; Lesenfants et al., 2019] was recently developed and has been shown to be measurable in CI users [Somers et al., 2018; Verschueren et al., 2019]. It is based on neural tracking of the speech envelope and can be measured from single-trial EEG recordings while listening to natural speech [Ding and Simon, 2013; Vanthornhout et al., 2018]. Since restoring speech understanding is one of the most important goals of cochlear implantation, it is likely a well-suited diagnostic to drive automated fitting algorithms. Genetic algorithms have been proposed to explore the large subject-dependent parameter space involved in fitting CIs [Durant et al., 2004; Wakefield et al., 2005].

Another application of EEG responses to continuous speech is the problem of auditory attention detection (AAD). Different speech streams are represented as separate auditory objects in the brain [Shinn-Cunningham, 2008] and the representation of the attended speaker is stronger [O’Sullivan et al., 2015]. Machine learning techniques have been successfully applied to EEG recordings to determine to which of multiple speakers the listener is attending, which can be used to steer the directional gain and noise suppression in so-called neuro-steered hearing devices [Van Eyndhoven et al., 2016; Das et al., 2018; Geirnaert et al., 2019; Aroudi and Doclo, 2020]. While most of the published research so far has focused on neuro-steered hearing aids, it has been shown that AAD in CI users is feasible [Nogueira et al., 2019; Paul et al., 2020].

In conclusion, enabling cochlear implants to record EEG signals and process them into relevant auditory diagnostics is a prerequisite for the development of new applications that can better rehabilitate deaf or severely hearing-impaired persons. In this study, we demonstrated the usage of CI electrodes to record and characterise continuous EEG, and the capability of obtaining objective auditory diagnostics from these recordings. This paves the way towards chronic neuro-monitoring of hearing-impaired users in their everyday listening environment, and neuro-steered hearing prostheses.

## IV. Methods

### A. Participants

Three adult CI users with a percutaneous connector on their implant participated in this study. The subjects took part in a research program by Cochlear Ltd, during which the subjects were implanted with an experimental cochlear implant that has an percutaneous connector, providing direct access to the implanted electrode leads (see section IV-B). Due to this experimental implant, these subjects are rare. Because only three subjects were enrolled in the research program at the time of this study, the sample size of our study is limited. The subjects took part in the research program for two years, after which the percutaneous device is explanted and replaced with a commercial CI. Subjects were carefully screened and briefed before enrolment in the program, and signed an informed consent approved by Western IRB. The subjects were free to terminate their participation at any point during the program without repercussions. Relevant individual subject details are shown in Table I.

**TABLE I.** Subject details

| Subject | Age (years) | Gender | CI ear | Date tested | Gain | Stimulated electrodes |
| --- | --- | --- | --- | --- | --- | --- |
| S1 | 30 | F | L | Nov. 2018 | 10000 | E21-E13 |
| S2 | 56 | F | R | May 2019 | 20000 | E04-E12 |
| S3 | 48 | F | L | May 2019 | 20000 | E04-E12 |

### B. Percutaneous cochlear implant

The cochlear implant with percutaneous connector used in this study is functionally equivalent to commercial Cochlear devices, however the wireless radio frequency (RF) link between implant and behind-the-ear piece is replaced by a wired connection through the skin. The implanted part consists of a conventional 22-contact intracochlear Nucleus® Contour Softip(tm) perimodiolar electrode array, and two extracochlear reference electrodes located below the temporalis muscle. The extracochlear electrodes are referred to as MP1 and MP2 (i.e. monopolar electrodes 1 and 2). The MP1 electrode is a small ball electrode placed. In conventional CIs, the MP2 electrode is a plate electrode attached to the casing of the implant. As there is no implant casing present with the percutaneous device, the MP2 electrode is a flat plate electrode placed under the temporal muscle. The intracochlear electrodes are numbered E01 to E22, counting from base towards the apex of the cochlea.

All implanted electrode leads are terminated to a connector housed in a percutaneous titanium pedestal mounted to the skull behind the ear. For take-home use, each subject received a behind-the-ear piece with an adaptor that fits the percutaneous connector. In addition to a microphone, sound processor and battery, this external piece was equipped with a stimulator that is electrically equivalent to a Nucleus® Contour Advance(tm) implant. For the purposes of this study, an electrode access board is connected to the percutaneous connector with a cable, making the electrode leads easily accessible for external equipment. The external electrical stimulator as well as the recording amplifiers are connected to pins of the electrode access board during the experiments. The stimulation and recording devices can be connected to leads of any of the 22 intracochlear electrode or the 2 extracochlear reference electrodes.

### C. Experimental set-up

A schematic representation of the experimental set-up is depicted in Fig. 7. Stimulation and recording are controlled from a battery-operated laptop to avoid any connections to mains voltage.

**Fig. 7.**
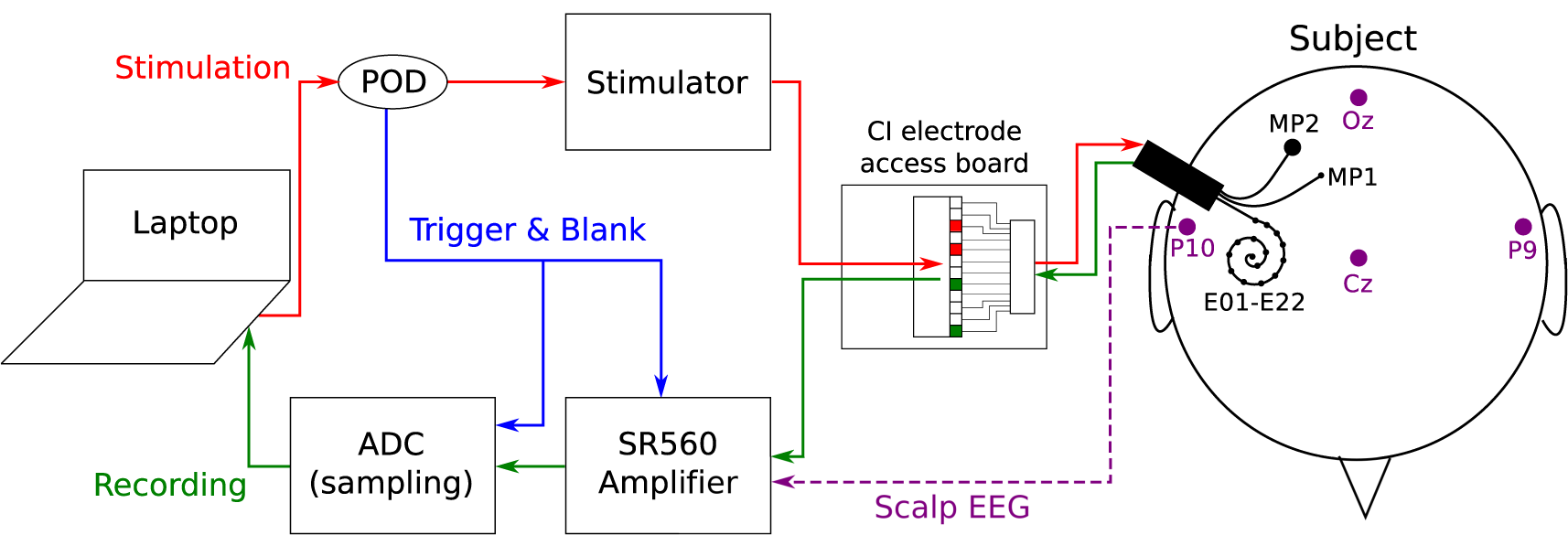
Experimental set-up. The electrical stimulation path is shown in red, and the intracochlear EEG recording path is shown in green. Additional scalp electrodes are placed on the subject’s head, shown in purple. The trigger signal used to synchronize EEG recordings and control the amplifier blanking is shown in blue.

*1) Stimulation path:* The electrical stimulation is generated through a custom software platform developed at the ExpORL research lab [Hofmann and Wouters, 2010] that allows interfacing with Cochlear’s Nucleus® devices through the Nucleus Implant Communicator (NIC). The generated sequences are sent to a programming device (POD) connected to an SP12 speech processor. Conventionally, an SP12 is worn behind the ear and transmits the processed sounds to the implant through a RF link. We used an SP12 with modified adaptor. Instead of an RF transmission coil, it was fitted with a connector interfacing with the electrode leads on the electrode access board.

*2) Recording path:* Two SR560 low noise, single-channel, analog voltage preamplifiers (Stanford Research Systems) were used to amplify the voltage signals on the implanted electrodes. Each SR560 was connected to two electrode leads on the access board, i.e. an active and reference electrode for the EEG signal. The amplifiers were configured to amplify in differential mode to reduce the effects of noise on the electrode leads. Unless specified otherwise, an additional electrode on the subject’s collarbone was used as the ground for the differential recordings. The analog outputs of the SR560 were sampled using an NI USB-6216 data card (National Instruments) as an Analog-to-Digital Converter (ADC). The digitized samples are transferred to the laptop through USB and stored for later offline analysis.

In some trials, scalp electrodes were used to record EEG signals for comparison with the implanted electrode recordings. In this case, the input of the SR560 amplifier was changed to a scalp electrode lead instead of a lead on the electrode access board. The recording settings of the SR560’s and the rest of the recording path remained unchanged when using scalp electrodes. Either the active lead, reference lead, or both, could be changed to a scalp electrode instead of a CI electrode. This allows flexibility to perform recordings between two scalp electrodes, between two implanted electrodes, or hybrid recordings between an implanted and a scalp electrode (see section IV-F).

The ADC has a maximal total sampling rate of 400 kHz across all channels. Thus, the three channels (2 EEG signals plus the trigger signal, see below) are each recorded at 133 kHz. An analog anti-aliasing filter with cut-off 30 kHz with 6dB/octave roll-off is enabled on both SR560s. The amplifier gains are adjusted to the maximal possible values before signal clipping occurred, dependent on the subject (see Table I).

*3) Control and safety circuitry:* The stimulating POD was configured to emit a trigger pulse on every stimulation pulse. This trigger signal serves two purposes. Firstly, it is recorded by the ADC alongside the EEG signals from the amplifiers. The recorded trigger signal is used for synchronization and for cutting the EEG into trials during the offline analysis. Secondly, the trigger signal is connected to the blanking input of the SR560s. While this input is high, the amplifier gain is set to zero, i.e. the recorded signals are “blanked” during the triggers. This ensures that the large electrical stimulation artifacts are not amplified, which may cause the amplifier to saturate. The blanking length is controlled with a custom-made circuit that changes the duration of the trigger pulse at the blanking input. Additional measures are taken to ensure the subject’s safety. The entire set-up is DC powered by batteries, to ensure that there are no possible connections with mains voltage. A DC blocking circuit is inserted between the amplifier input and the access board to prevent any leakage currents of the amplifier’s input transistors from flowing back to the subject. Isolation barriers are present at several locations in the set-up to create galvanic separations between the subject and any equipment that may fault.

### D. Stimuli

The targeted EABR and ECEP responses are evoked using biphasic electrical click trains. The pulses are presented in bipolar configuration, i.e. the stimulating voltage is applied between two intracochlear electrodes. The advantage of using a bipolar stimulus is that both extracochlear reference electrodes MP1 and MP2 are available for recordings. For every subject, the electrodes with the lowest thresholds are selected as the stimulation electrodes, and are shown in Table I. The pulse width of each phase is 100 µs with an inter-phase gap of 8 µs. The polarity of the pulses is alternated every trial in order to reduce the stimulation artifact when averaging.

For the EABR, a single biphasic pulse is repeated at 27 Hz. In total, 3000 trials are recorded, resulting in a recording time of 2 minutes per EABR trace. The blanking length is set to 1 ms. For the ECEP, a 2000 Hz burst of biphasic pulses sustained for 5 ms is repeated at 1 Hz. In total, 300 trials are recorded, resulting in a recording time of 5 minutes for a single ECEP trace. The blanking length is set to 10 ms such that the artifact of the entire pulse burst is blanked. To prevent the implant from losing power, power-up frames are inserted in the stimulus sequences every 1 ms, except for the 12 ms post-stimulus interval. These frames only deliver power to the implant and do not stimulate the neural tissue. They cause a small electrical artifact which is removed through linear interpolation (Fig. 8A).

**Fig. 8.**
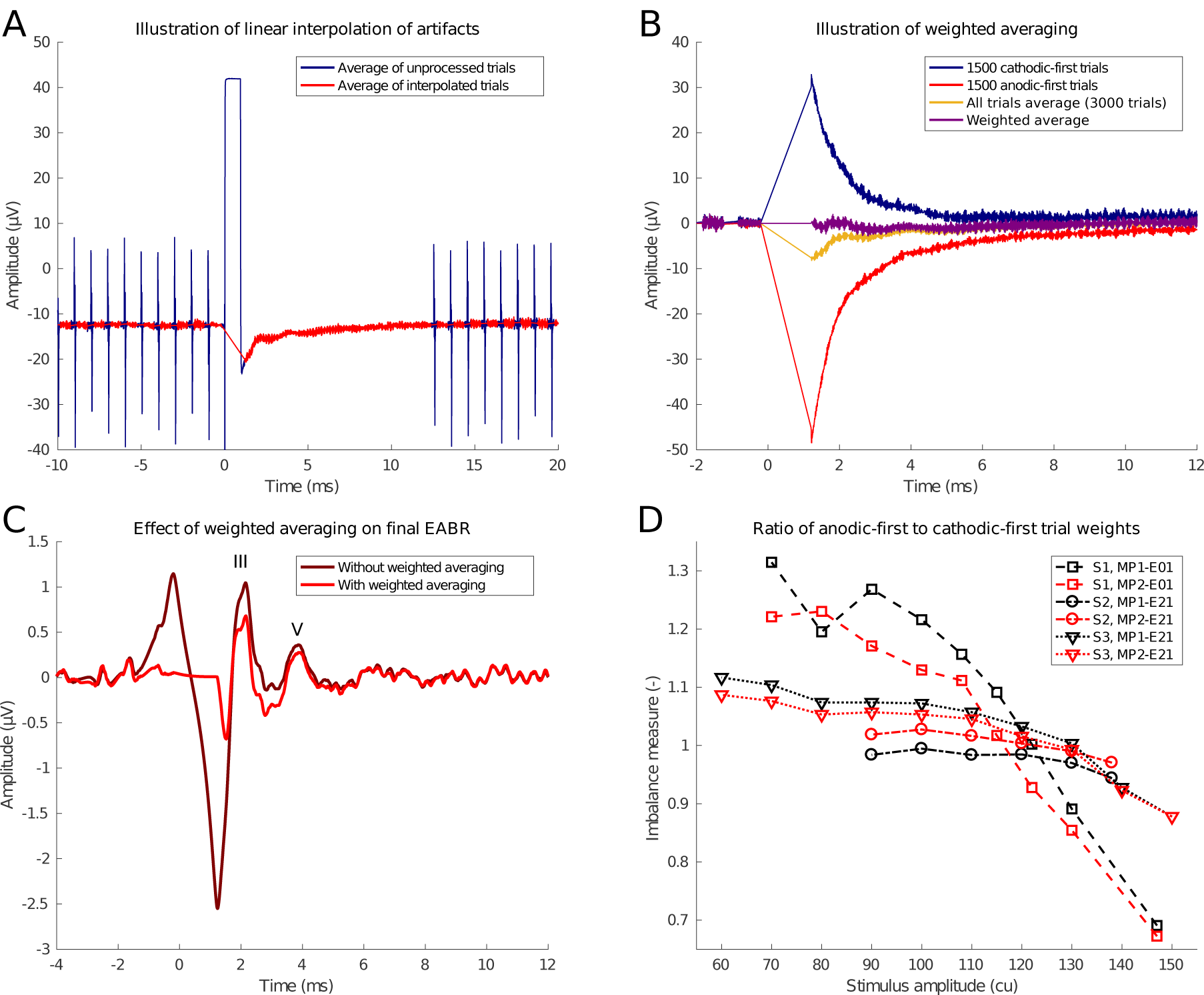
Illustration of signal processing techniques. This is an example from an EABR at C-level from subject S1, recorded between electrodes MP2-E01 using bipolar stimulation between electrodes E21-E13. **A** Example of pedestal artifact caused by amplifier blanking, and power-up frame artifacts before and 12 ms post-stimulus. These artifacts are removed by linear interpolation. **B** Example of weighted averaging procedure to remove sawtooth artifact introduced by linear interpolation. **C** Final EABR trace after signal processing with and without weighted averaging. Waves III and V are marked. **D** Imbalance between amplitudes of exponentially decaying artifacts as function of stimulation level. The imbalance measure is the ratio of the weight assigned to the anodic-first trials divided by the weight assigned to the cathodic-first trials. Values greater than 1 indicate that anodic-first trials receive a greater weight, i.e. the cathodic-first trials exhibit a larger stimulation artifact, and vice versa for values smaller than 1.

### E. Signal processing and Artifact removal

Every stimulation pulse causes a large stimulation artifact in the EEG recording. Especially as stimulation and recording electrodes are both located within the cochlea, the artifacts were hypothesized to be extremely large. The artifact consists of two main components: a large, biphasic artifact corresponding to the biphasic stimulation pulse, and a subsequent transient exponential decay, caused by the remaining charge imbalances redistributing in the cochlea. The amplifier blanking eliminates the former part of the stimulus artifact completely. However, during blanking, a pedestal artifact appears (Fig. 8A). Even though the gain is zero during blanking, small differences between amplifier and subject grounds cause this pedestal artifact. This artifact, along with the artifacts from the power-up frames, can be easily removed using linear interpolation between a sample before and a sample after the artifact [Hofmann and Wouters, 2010], as depicted in Fig. 8A.

The remaining exponentially decaying tail of the artifact can last several milliseconds. In the case of EABRs, the tail coincides with the targeted evoked response, so it cannot be removed by blanking. Furthermore, the end point of the linear interpolation is on the exponential tail, which produces a triangular, sawtooth-like artifact (Fig. 8B). The polarity of the exponential tail depends on the polarity of the stimulus, whereas the response only differs minimally for different polarities. Since the trials were presented with alternating polarities, the sawtooth artifact can be reduced by averaging both polarities together. Because of non-linearities in the electrode-tissue interface, the artifacts caused by both polarities are not symmetric. Using a weighted averaging procedure which assigns lower weights to the polarity with the largest artifact amplitude [Alvarez et al., 2007], the residual sawtooth artifact can be eliminated completely (Fig. 8B). This weighted averaging procedure is employed in all analyses in this study. Fig. 8C shows an example of what happens with the EABR response when the sawtooth artifact is not corrected: due to filtering the artifact smears out and distorts the EABR waves. This is undesirable as the sawtooth artifact amplitude depends on stimulus intensity and may lead to inflated or false positive EABR responses. Interestingly, the imbalance between artifact amplitudes in anodic-first and cathodic-first trials depends on the stimulus intensity (Fig. 8D), i.e. the weighted averaging procedure requires different weight ratios depending on stimulus intensity. This indicates that the effect of the electrode-tissue interface on the stimulus artifacts is non-linear, and depends strongly on the impedances and capacitive characteristics of the surrounding tissue [Kent et al., 2015; Freche et al., 2018].

In addition to the aforementioned steps to eliminate stimulus artifacts, some other preprocessing is applied before visualizing the EEG responses. A baseline correction is applied to remove signal drift using a pre-stimulus interval as the baseline (EABR: 5 ms, ECEP: 100 ms). All trials exceeding a certain threshold were rejected from the analysis (EABR: 25 µV, ECEP: 50 µV). This rejects trials containing other non-stimulus related artifacts, such as amplifier clipping or biological artifacts. Remaining trials are filtered within a specific passband suited for the evoked response (EABR: 300 Hz to 3000 Hz, ECEP: 1 Hz to 30 Hz).

### F. Experiments

Before any EEG recordings were performed, appropriate stimulation levels were determined for all subjects using a behavioural method. For every stimulus, the intensity was gradually increased from zero until the subject indicated that a loud but comfortable level was reached, which was marked as the comfortable (C) level. Next, the intensity was decreased again until the minimum intensity which caused an audible percept was reached, marked as the threshold (T) level. C and T levels were determined for every stimulus. Any change in the stimulus (e.g. repetition rate, pulse width, stimulating electrode) requires a new C and T-level determination. Before connecting the electrode access board leads or scalp electrodes to the recording amplifiers, the impedances of the electrodes were checked to be within an acceptable range (scalp: *<* 5 kΩ, implanted: *<* 10 kΩ).

The first experiment conducted in every participant was to measure an EABR intensity sweep to validate the set-up and signal processing. We chose this paradigm to ensure that responses were detectable, not influenced by artifacts, and exhibited a normal amplitude growth. The sweep consisted of recording EABR responses to stimuli with intensities ranging from the subject’s C-level to below T-level. The presentation order of different intensities was randomized. EABR responses were recorded from electrodes MP1-to-Exx and electrodes MP2-to-Exx, where Exx is an intracochlear electrode chosen depending on the subject.

An important goal of this study is to identify the optimal electrode configuration to record evoked potentials using implanted electrodes. Because MP1 and MP2 are implanted and cannot be moved, we made use of scalp electrodes to obtain “hybrid” implanted electrode recordings, i.e. between an implanted electrode and a scalp electrode. The scalp electrode can be freely placed anywhere on the head, and as such can be used to simulate an additional implanted electrode near that site. For both the EABR and ECEP responses, recordings with different electrode configurations were performed and compared. All of these recordings used C-level stimulation. Scalp electrodes were placed on the head in locations commonly used in evoked response studies: left/right mastoid (P9/P10), top of the head (Cz), back of the head (Oz) (see Fig. 1). In addition to these hybrid recordings, recordings with only implanted or only scalp electrodes were performed.

## Acknowledgements

This project has received funding from the European Research Council (ERC) under the European Unions Horizon 2020 research and innovation program (grant agreement No. 637424, ERC starting grant to Tom Francart). Ben Somers is supported by a PhD grant for Strategic Basic research by the Research Foundation Flanders (FWO, No. 1S46117N). We thank Cochlear Ltd. for providing the necessary research equipment and testing time with the CI users with percutaneous connector.

Fig. 6 is composed with icons from thenounproject.com, with following titles and artists: “Cochlear Implant” by Ben Davis, “Brain” by David, “point” by Chameleon Design, “Choice” by Mat Fine, “Activity” by mikicon.

## Author contributions statement

B.S. and T.F. contributed to the design and conception of the study. B.S. and C.J.L. contributed to the design of the experimental set-up and conducted the experiments. B.S analysed the data and wrote the manuscript. All authors reviewed and edited the manuscript.

